# Synthetic Longitudinal Tabular Data Generation via Copula

**DOI:** 10.64898/2026.08.03.742474

**Authors:** Hanchang Cai, Wenshan Yu, Ruijin Lu, Ishanu Chattopadhyay, Xinlian Zhang, Jinyuan Liu

**Affiliations:** Division of Biostatistics and Bioinformatics, Herbert Wertheim School of Public Health and Human Longevity Science, University of California San Diego, 9500 Gilman Drive, La Jolla, CA 92093, United States; Department of Population Health Sciences, School of Public Health, Georgia State University, 140 Decatur Street, Atlanta, GA 30303, United States; Institute for Informatics, Data Science & Biostatistics, Washington University in St. Louis School of Medicine, 660 S. Euclid Ave., St. Louis, MO 63110, United States; Institute for Biomedical Informatics, Department of Internal Medicine, University of Kentucky College of Medicine, 800 Rose Street, Lexington, KY 40536, United States; Departments of Biostatistics, Vanderbilt University Medical Center, 2525 West End Avenue, Suite 1100, Nashville, TN 37203, United States

**Keywords:** generative adversarial network (GAN), missing data, multiple imputation, synthesis variability, synthetic data evaluation, temporal correlation in longitudinal data

## Abstract

Synthetic data generation is increasingly used to enable data sharing and secondary analysis while protecting participant privacy, particularly for longitudinal tabular health data, where repeated measures per subject create within-subject dependence that most synthetic data methods are not designed to preserve. Existing generative methods, particularly generative adversarial network (GAN)-based approaches, can model complex distributions, but their estimated dependence structures are often difficult to interpret and their performance may be unstable or prone to overfitting in modestly sized datasets. Here we show that eCDF-copula, a statistically rooted approach using the empirical cumulative distribution function (eCDF) and copula modeling, preserves within- and between-visit dependence structure. To handle pervasive missing data, we propose a two-stage strategy combining multiple imputation with copula-based synthesis, enabling a variance decomposition that quantifies replication variability across methods. We benchmarked the proposed approach against four established methods on two longitudinal clinical datasets spanning markedly different sample sizes (*n* = 120 vs. *n* = 3, 612). eCDF-copula achieved resemblance and utility exceeding those of state-of-the-art synthetic data methods, while maintaining comparable privacy.

## 1. Introduction

The increasing use of electronic health records (EHR), clinical registries, and large-scale observational studies has created unprecedented opportunities for developing predictive models, evaluating treatment effectiveness, and advancing precision medicine (Yale et al., 2020). However, access to these data is often restricted by privacy regulations, institutional governance policies, and persistent underrepresentation of important patient subgroups (Lewis et al., 2023). Synthetic data, artificial observations generated by a statistical or machine-learning model trained to approximate the joint distribution of real data, have therefore emerged as a means of expanding data accessibility while preserving the statistical characteristics of the original data (Hernadez et al., 2023). Beyond supporting privacy-preserving data sharing and methodological development, synthetic data also enables augmentation of limited datasets for predictive modeling (Alqahtani et al., 2021). Within biomedical research, synthetic data generation has been explored across modalities including physiological signals, medical imaging, free-text electronic health records, and activity monitoring (Yan et al., 2022), yet structured tabular data is still among the most extensively used, given its central role in clinical prediction, causal inference, and decision-support analyses.

However, most current synthetic data methods, including generative adversarial networks (GANs) and related deep generative approaches (Gulrajani et al., 2017), function as black boxes, offering limited statistical guarantees (Xu et al., 2019). The reliance on large training datasets also makes them ill-suited to the small-to-moderate sample sizes that characterize most clinical studies (Heine et al., 2023), including randomized controlled trials (RCTs). Even statistically grounded methods rarely account for longitudinal dependence, a limitation compounded by the missing data common to repeated clinical assessments (Rubin, 1987). These shortcomings are further obscured by a lack of comprehensive fidelity benchmarking: empirical evaluation spanning *resemblance, utility*, and *privacy* is rarely undertaken in full prior to the dissemination (Hernadez et al., 2023; Snoke et al., 2018). Consequently, statistically transparent, comprehensively validated approaches for generating longitudinal synthetic tabular data that faithfully preserve the dependence structures required for reliable downstream analysis and data sharing remain lacking.

In this paper, we address these gaps by developing a synthetic data generation method for longitudinal tabular data that is transparent, statistically grounded, and broadly applicable to the moderate-sized datasets typical of clinical research. Rather than relying on black-box methods, our approach leverages the probability integral transform (van der Vaart, 1998) and copula functions (Lambert and Vandenhende, 2002), two classical statistical constructs with well-established theoretical properties, to separate marginal distributions from dependence structure, thereby preserving interpretability and statistical accountability that deep generative methods do not provide. This framework extends naturally to longitudinal data, capturing dependence across repeated measures while accommodating the missing data inherent to clinical follow-up (Hamori et al., 2020). Using two real-world longitudinal datasets, we evaluate this method against off-the-shelf alternatives, including Conditional Tabular GAN (CTGAN), Wasserstein GAN with Gradient Penalty (WGAN-GP), Gaussian multivariate (GM), and probabilistic autoregressive model (PAR), through comprehensive benchmarking spanning resemblance, utility, privacy (El Emam et al., 2022), and synthesis variability.

The remainder of the paper is organized as follows. Section 2 describes the eCDF-copula method for cross-sectional and longitudinal data, along with the two-stage strategy for handling missing data. Section 3 details the study datasets, competing methods, and evaluation framework. Section 4 presents results across resemblance, utility, privacy, and synthesis variability. We conclude in Section 5.

## 2. Methods

### 2.1. Background

#### 2.1.1. Empirical Marginals

Let *Z* be a continuous random variable with cumulative distribution function (CDF) *F* (*z*). The probability integral transform states that *U* := *F* (*Z*) ~ Unif(0, 1) follows a Uniform(0,1) distribution, and conversely, *Z* = *F* ^*−*1^(*U*) follows the distribution *F* for any *U* ~ Unif(0, 1) (Casella and Berger, 2002). This inverse transform relationship underlies the distribution-agnostic simulation proposed in this paper.

In biomedical applications, the true CDF, or underlying distribution, *F* (*z*) is typically unknown and often highly irregular, e.g., by exhibiting zero inflation, heavy tails, or multimodality (Alqahtani et al., 2021). We therefore substitute the empirical CDF (with *I*(·) an indicator function):

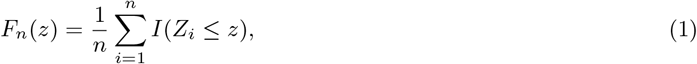

whose generalized inverse, 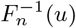, maps uniform draws to the quantiles of observed data. This substitution preserves marginal features (e.g., skewness, heavy tails, bounded support) without imposing any parametric distribution, and forms the “e” in the proposed eCDF-copula approach.

#### 2.1.2. Copula and Dependence Modeling

Let **Z** = (*Z*_1_, *Z*_2_, …, *Z*_*p*_)^*⊤*^ be a *p*-dimensional random vector with joint CDF *F*_**Z**_(*z*_1_, *z*_2_, …, *z*_*p*_) and continuousmarginal CDFs (*F*_1_, *F*_2_, …, *F*_*p*_). A *copula* is a multivariate CDF on the unit cube [0, 1]^*p*^ with uniform univariate marginals (Nelsen, 2006), and by Sklar’s theorem, there exists a unique copula *C*(·) linking the joint distribution to the marginals through the uniform-transformed variables *F*_1_(*z*_1_), …, *F*_*p*_(*z*_*p*_) (Sklar, 1959):

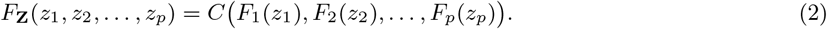

Therefore, copulas can separate marginal distributions from their dependence structure. We use this decomposition for synthetic data generation: empirical marginals preserve each variable’s distribution, while a fitted copula preserves the dependence structure among variables.

#### 2.1.3. Gaussian Copula and Rank Estimation

We focus on the Gaussian copula parameterized by a correlation matrix **R**,

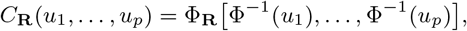

where ϕ_**R**_(·) is the *p*-variate normal CDF and ϕ^*−*1^(·) is the standard normal quantile function. This choice is motivated by computational practicality and empirical performance: it requires estimating only a single *p* × *p* correlation matrix, admits robust rank-based estimation without parametric marginal assumptions, and remains computationally stable in moderate-to high-dimensional settings, as shown in our benchmarking comparison. Other copula families, including *t*, Archimedean, and extreme-value copulas (Joe, 1997), can be substituted within the same framework when tail dependence or asymmetric dependence structures are of interest.

Because copulas are invariant under strictly monotone marginal transformations (Nelsen, 2006), the dependence parameters in **R** can be estimated from ranks rather than from the original measurement scales. We use Spearman’s correlation *ρ*_*s*_, defined as the Pearson correlation between the probability-transformed variables *F*_*j*_(*z*_*j*_) and *F*_*j*′_ (*z*_*j*′_) for variable pairs *j, j*′ = 1, …, *p*. For the Gaussian copula, Spearman’s correlation is related to the latent Gaussian correlation by *ρ*_*s*_ = (6*/π*) arcsin(*R*_*jj*_*′ /*2) (Kruskal, 1958), so *R*_*jj*′_ can be estimated by inverting this relationship. In our implementation, we use the sample Spearman correlation matrix as an approximation. For the low-to-moderate correlations encountered in many applications, the resulting numerical difference is small (Liu et al., 2009).

### 2.2. Cross-sectional Synthetic Data Generation

Building on the statistical ingredients introduced above, we now assemble the proposed framework into a cross-sectional synthesis algorithm. Consider a tabular dataset **Z**^(*n×p*)^ = {*Z*_*ij*_}, *i* = 1, …, *n, j* = 1, …, *p*, where rows index subjects and columns index variables. In this setting, each subject contributes a single *p*-dimensional vector, and the goal is to generate a synthetic dataset 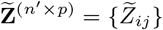 with user-defined sample size *n*′, typically chosen so that *n*′ ≥ *n*, while preserving the marginal distributions and between-variable dependence in **Z**. This algorithm also provides the building block for the longitudinal extension in Section 2.3.

Generating *n*′ ≥ *n* synthetic records is often useful in practice, especially for training flexible machine learning models at a larger scale or for class rebalancing in imbalanced settings (Restrepo et al., 2023; Ben Hassine and Mili, 2025). We proceed in four steps, summarized in **Algorithm 1**.

#### Step 1: Empirical Marginals

For each variable *j*, we estimate the marginal distribution using the empirical CDF as in (1):

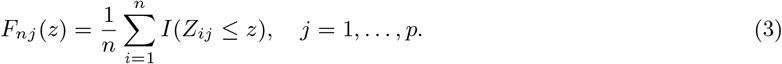

After applying a prespecified rule for handling ties, such as average ranks or random tie-breaking in standard software (R Core Team, 2026), we compute the scaled ranks

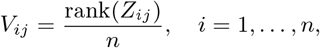

which serve as pseudo-observations for variable *j*.

#### Step 2: Rank-based Dependence Estimation

Let **Z**_*·j*_ = (*Z*_1*j*_, …, *Z*_*nj*_)^*⊤*^ denote the observations for variable *j*, and let **V**_*·j*_ = (*V*_1*j*_, …, *V*_*nj*_)^*⊤*^ denote its pseudo-observations from Step 1. We estimate the dependence structure by computing the *p* ×*p* sample Spearman correlation matrix **R**, with entries

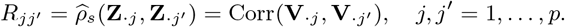

If **R** is not positive definite (PD), we can project it to the nearest positive definite correlation matrix.

#### Step 3: Copula Simulation

Next, we draw *n*′ independent samples from the Gaussian copula with correlation matrix **R** by sampling **W**^(*i*)^ ~*N*_*p*_(**0, R**) and setting *U*_*ij*_ := ϕ(*W*_*ij*_), yielding

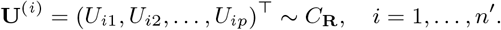

This step encodes the dependence structure through copula and marginally uniform variables *U*_*ij*_.

#### Step 4: Inverse Transform

Lastly, we map each simulated uniform variate *U*_*ij*_ back to the original data scale via

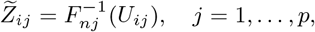

Where 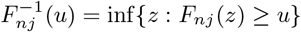 is the generalized inverse of the empirical CDF in (3). This inverse transform maps the copula-generated uniforms onto the observed scale of each variable *j*, so the resulting synthetic values follow its empirical marginal distribution and retain features such as the observed mean, variance, skewness, and tail behavior up to Monte Carlo variation. Moreover, since 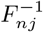 takes values on the support of variable *j*, the synthetic values 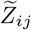 remain within that support without extrapolation.

By construction, the synthetic dataset 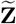 satisfies two fidelity guarantees. First, the marginal distribution of 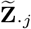_*·j*_ converges to *F*_*j*_ as *n* → ∞, since *F*_*nj*_ → *F*_*j*_ uniformly almost surely by the Glivenko-Cantelli theorem, and 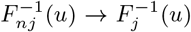 at all continuity points of 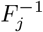 (van der Vaart, 1998). Second, when correlations are low to moderate, the pairwise Spearman correlations of 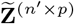 approximate those of **Z**^(*n×p*)^ well (Hoff, 2007; Liu et al., 2009).

### 2.3. Longitudinal Synthetic Data Generation

Longitudinal studies repeatedly measure *p* variables across *T* visits, which induce two types of dependence: variables measured at the same visit tend to correlate with one another (within-visit dependence, as in the prior cross-sectional setting), and measurements from the same subject may be correlated across visits (between-visit temporal dependence) (Diggle et al., 2002). An effective generator should preserve both within-visit and between-visit dependence.

Let the observed dataset be **Z**^(*n×p×T*)^ = {*Z*_*ijt*_}, where *i* = 1, …, *n* indexes subjects, *j* = 1, …, *p* indexes variables, and *t* = 1, …, *T* indexes visits. The goal is to generate a synthetic dataset 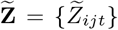 with user-defined sample size *n*′, typically *n*′ ≥ *n*, while preserving visit-specific marginal distributions and the joint dependence structure across variables and visits.

#### Algorithm 1

Cross-Sectional eCDF-Copula Synthesis

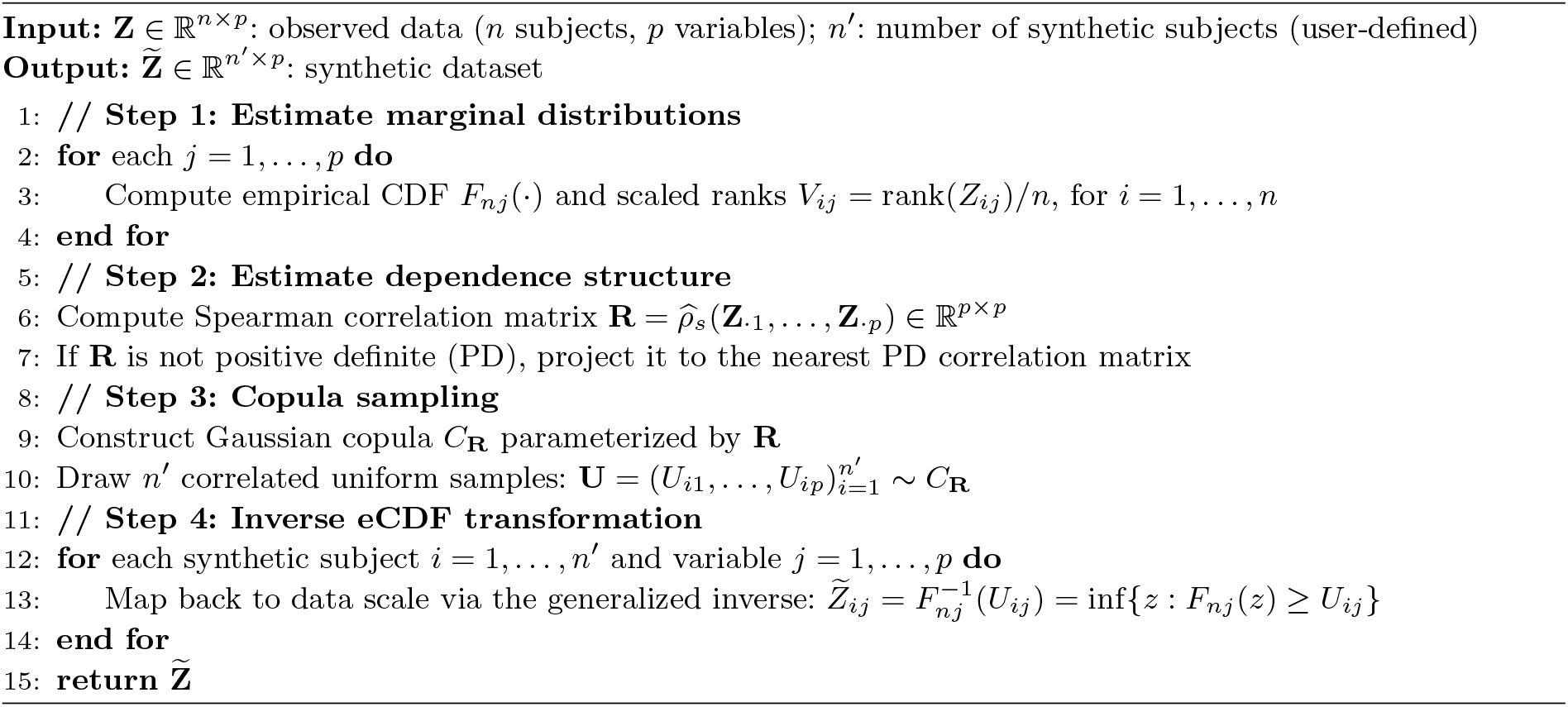

#### Wide-format Representation

We first arrange the longitudinal data in wide format by concatenating the visit-specific matrices column-wise. Let **Z**_*t*_ = (*Z*_*ijt*_)_*i*=1,…,*n*; *j*=1,…,*p*_ ∈ ℝ^*n×p*^ denote the matrix of observations at visit *t*. We define

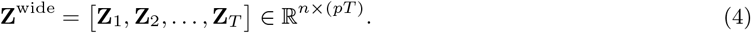

Thus, each row of **Z**^wide^ contains all variables for one subject across all visits. This representation converts the longitudinal synthesis problem into a Gaussian copula problem on (*pT*) variables, to which the cross-sectional procedure in Section 2.2 can be applied.

#### Dependence Structure

Because variables are ordered by visit in **Z**^wide^, the Gaussian copula correlation matrix **R** ∈ ℝ^*pT ×pT*^ has a natural block structure:

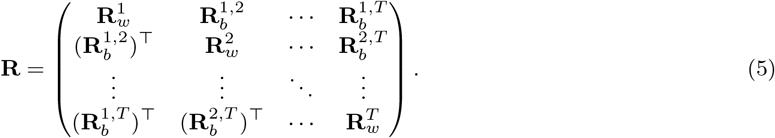

The diagonal blocks 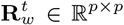 capture within-visit dependence among variables at visit *t*, whereas the off-diagonal blocks 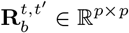 capture cross-visit dependence between variables measured at visits *t* and *t*′.

#### Synthetic Data Generation

We draw *n*′ independent samples from the Gaussian copula with correlation matrix **R** by sampling

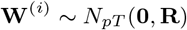

and setting

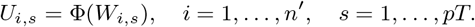

For each stacked variable *s*, we apply the empirical inverse CDF:

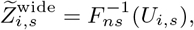

where *F*_*ns*_(·) is the empirical CDF of the *s*-th column of **Z**^wide^. Finally, we reshape 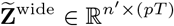 back into visit-specific matrices 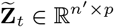 using the index mapping

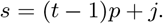

By construction, the synthetic data preserve visit-specific marginal distributions through the empirical inverse CDFs and reproduce both within-visit and between-visit dependence through the block-structured copula correlation matrix in (5).

In the applications considered here, we estimate the full block-structured correlation matrix **R** from the wide representation. When *p* and *T* are large relative to *n*, however, estimating all *O*(*p*^2^*T* ^2^) entries may be unstable (Mieldzioc et al., 2021). In such settings, the same framework can accommodate parsimonious block structures, for example, by replacing each within-visit block with an exchangeable correlation matrix and each between-visit block with a scalar multiple of the identity matrix:

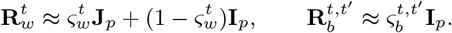

Here, **I**_*p*_ is the *p* × *p* identity matrix and **J**_*p*_ is the *p* × *p* matrix of ones. The scalar 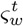 denotes the common within visit correlation among variables at visit *t*, and 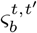 denotes the common within-variable temporal correlation between visits *t* and *t*′. This reduces the number of dependence parameters from *O*(*p*^2^*T* ^2^) to *O*(*T* ^2^), at the cost of assuming exchangeable within-visit correlations and common temporal correlations across variables.

#### Synthesis-induced Uncertainty

Let 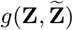 denote a pre-specified evaluation metric comparing the real dataset **Z** with a synthetic dataset 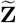. In this paper, *g*(·, ·) may represent any resemblance, utility, or privacy metric described in Section 3.3, such as a marginal distribution distance. The replication framework below should be applied separately to each metric of interest.

In the absence of missing data, variation in 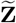 arises from stochastic copula sampling. To quantify this synthesis-induced uncertainty, we generate *L* independent synthetic datasets 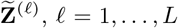, from the fitted generator and compute

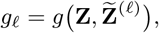

where the number of replicates *L* is chosen to stabilize the empirical mean and percentile range of *g*_*ℓ*_. We then summarize performance by the ensemble mean

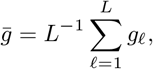

and characterize replication variability using the empirical standard deviation and the 2.5th–97.5th percentile range of 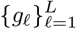

### 2.4 Handling Missing Data

Missing data are especially common in longitudinal studies, arising from participant dropout, missed visits, or intermittent non-response (Rubin, 1987). Unlike many deep-learning-based synthesis approaches, which embed missing-data handling within the generative model in ways that can be difficult to separate (Yoon et al., 2018; Mattei and Frellsen, 2019), we adopt a two-stage strategy that separates missing-data imputation from synthetic data generation. This separation allows variability due to imputation and variability due to stochastic synthesis to be quantified separately, thereby facilitating method comparison.

#### Stage 1: Multiple Imputation

Let **X**_*i*_ denote baseline and design covariates serving as auxiliary predictors, distinct from the longitudinal tabular variables to be synthesized. We work under the missing-at-random (MAR) assumption: conditional on the observed data and **X**_*i*_, the missingness mechanism does not depend on unobserved values. Including **X**_*i*_ as auxiliary predictors expands the conditioning set and can make the MAR assumption more plausible in practice (Collins et al., 2001).

##### Algorithm 2

Longitudinal eCDF-Copula Synthesis

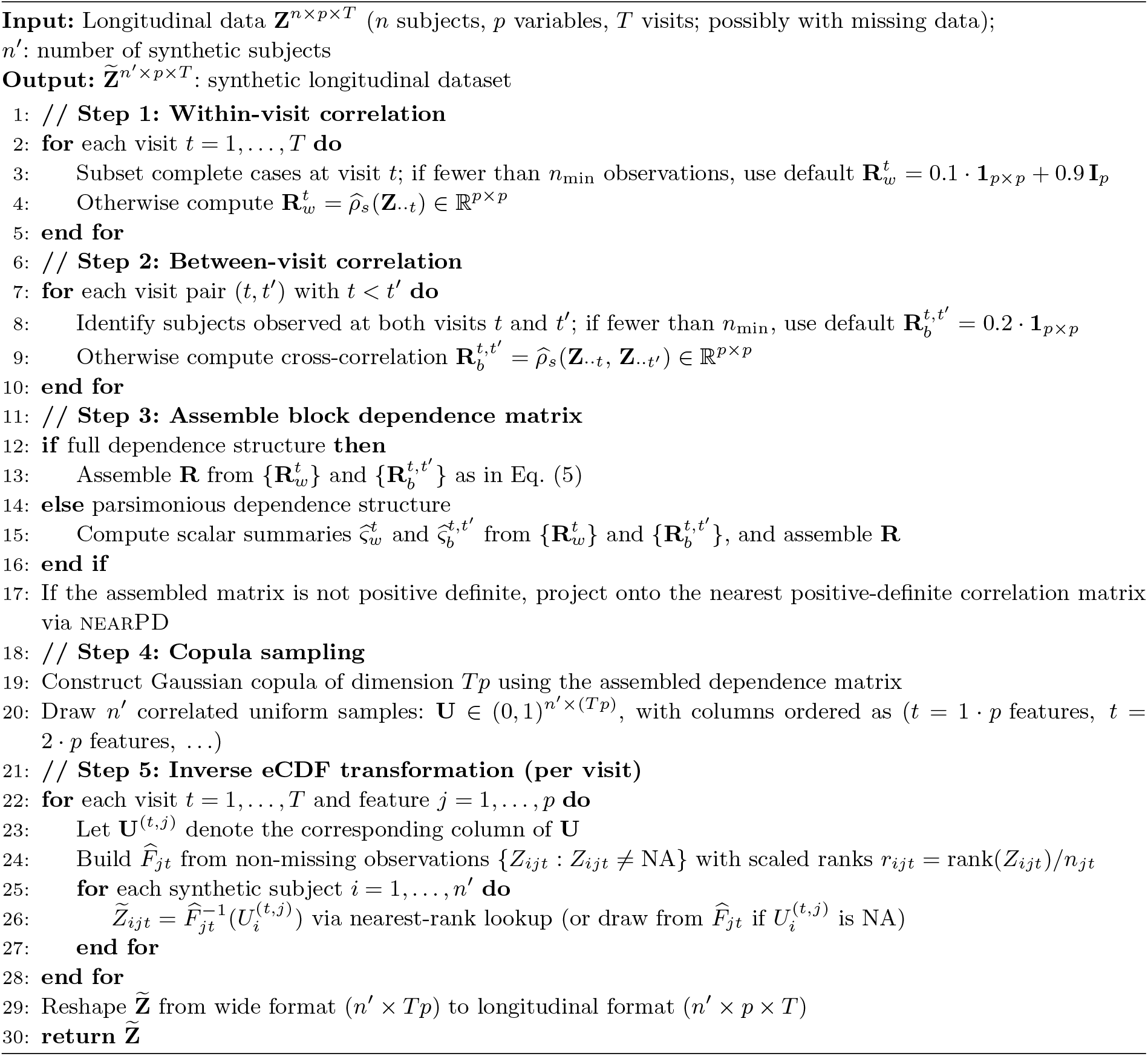

We handle missing data using predictive mean matching (PMM), as implemented in the mice package (van Buuren and Groothuis-Oudshoorn, 2011). Each subject’s *pT* variable-visit combinations are arranged in wide format, **Z**^wide^ ∈ R^*n×pT*^ as in (4), and augmented with auxiliary covariates **X** ∈ R^*n×q*^ before imputation. This wide-format imputation allows repeated measures and variables to inform one another without imposing a restrictive longitudinal correlation structure during imputation (Wijesuriya et al., 2025). For clustered or hierarchical data, the same framework can accommodate multilevel imputation models such as `2l.pan` or `2l.norm` (van Buuren, 2018). As in conventional multiple imputation, the procedure is repeated *M* times.

#### Stage 2: Synthesis and Variance Decomposition

For each of the *M* completed datasets, we apply the longitudinal eCDF-copula generator described in Section 2.3, drawing *L* synthetic datasets per imputation for a total ensemble of *M* × *L* synthetic datasets. As above, let *g*_*mℓ*_ denote a pre-specified scalar evaluation metric, applied separately to each resemblance, utility, or privacy measure of interest, computed on the *ℓ*-th synthetic dataset generated from the *m*-th completed dataset, where *m* = 1, …, *M* and *ℓ* = 1, …, *L*. The ensemble average is

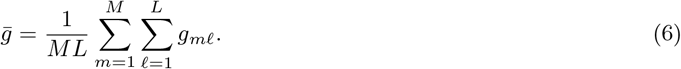

This nested design parallels multiple-imputation variance decomposition (Rubin, 1987; van Ginkel and Kroonenberg, 2014), but is used here as a descriptive diagnostic rather than for Rubin-style inference. It separates variability in the metric into contributions from the missing-data completion step and from stochastic synthesis conditional on a completed dataset. We summarize these sources using two empirical mean-square components:

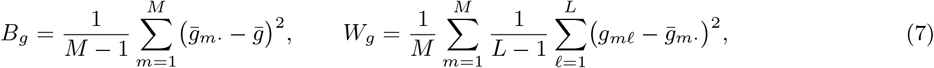

where 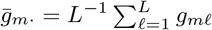 is the average metric value within the *m*-th completed dataset.

To interpret these components, suppose the metric follows

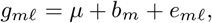

where *b*_*m*_ is the random effect for the *m*-th imputed dataset and *e*_*mℓ*_ is the synthesis-level deviation for the *ℓ*-th synthetic dataset generated from that imputation, with 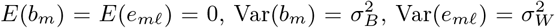 and *b*_*m*_ and *e*_*mℓ*_ independent.

Under this model, 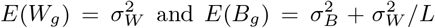. We therefore estimate the within-imputation *synthesis-induced uncertainty* and the between-imputation variance as

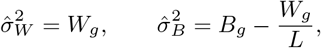

with negative between-imputation estimates truncated at zero if necessary. The total variability of the metric across synthetic datasets is then estimated by 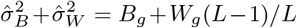 Although mean metric values remain the primary evidence for resemblance, utility, and privacy, this decomposition further assesses their stability across repeated imputation and synthesis. This is practically relevant when users analyze or release only one or a few synthetic datasets, so we included it as an additional evaluation dimension.

## 3. Study Design

### 3.1. Datasets

**Dataset A:** REMBRANDT is a multisite study of depression recurrence, cognitive burden, and neurobiological homeostasis in late life (Taylor et al., 2024). Participants were assessed at 14 scheduled visits over 24 months on depressive symptom severity (Montgomery-Åsberg Depression Rating Scale, MADRS) and anxiety severity (Hamilton Anxiety Rating Scale, HARS), yielding 28 longitudinal variables per participant. The sample comprises

*n* = 120 participants (40 Control, 80 Remitted), with a median of 13 observed visits (range 8–14); 42% had complete data across all visits, and the overall missing rate was 18% for both MADRS and HARS, concentrated at later visits due to dropout. Missing values were imputed via predictive mean matching (PMM), applied in wide format with demographic covariates as auxiliary predictors. Each feature was standardized to unit variance prior to synthesis.

**Dataset B:** The second dataset is from the Chronic Hypertension and Pregnancy (CHAP) trial (Tita et al., 2022), a longitudinal clinical study with repeated blood pressure measurements through delivery. We analyzed Period 2, comprising 3,934 pregnant participants (3,746 in Group 0; 188 in Group 1) with systolic (SBP) and diastolic (DBP) measurements between gestational weeks 3 and 39. Raw measurements were irregularly spaced (median 10 per participant, range 2–18), so we binned gestational weeks into ten consecutive 4-week intervals and averaged SBP and DBP within each bin per subject, yielding a wide-format matrix of 20 blood pressure features. Subjects missing more than 10 of these 20 features were excluded, corresponding to setting *n*_*min*_ = 10 in Algorithm 2, since fewer observed measurements may lead to unstable estimation of the correlation structure. This resulted in *n* = 3, 612 retained subjects, with a median of 14 observed features (range 12–20) and an overall missing rate of 26.8%.

Subjects were partitioned into 5 folds at the subject level, such that all visits from a given subject were assigned to the same fold; this partition was held fixed across all imputations. Missing values and synthesis then followed the two-stage procedure of Sections 2.4 within each fold, with *M* = 5 and *L* = 10, using synthetic data generated from the four training folds, and the held-out fold was reserved for downstream evaluations.

### 3.2. Competing Methods

We benchmarked the proposed eCDF-copula method against four established synthetic generators. For classical statistical paradigms, we considered:

- **Gaussian multivariate (GM)** constructs a multivariate distribution via a Gaussian copula fitted to parametric univariate marginals (Hernadez et al., 2023). This closest statistical comparator directly tests whether our nonparametric marginal eCDF component provides a meaningful benefit.
- **Probabilistic autoregressive model (PAR)**, implemented within the Synthetic Data Vault (SDV) ecosystem (Patki et al., 2016), is a longitudinal generative model that conditions on past observations to capture within-subject temporal dependence (Zhang et al., 2022).

For deep learning paradigms, we considered two variants of the generative adversarial network (GAN). Briefly, GANs learn to generate data through two networks trained in opposition: a *generator*, which produces synthetic observations from random noise, and a *discriminator*, which learns to separate real from generated observations while the generator learns to defeat it. Ideally, this adversarial process yields a generator that produces convincingly realistic data and a discriminator that has learned strong feature representations characteristic of the training data. However, Arjovsky et al. (2017) show that the Jensen-Shannon divergences typically minimized by GANs can be discontinuous with respect to the generator’s parameters, leading to training difficulty, and introduce the Wasserstein GAN (WGAN), which uses the Wasserstein loss to help stabilize training. A WGAN can still produce poor samples or fail to converge, since interactions between its weight constraint and the cost function can result in vanishing or exploding gradients. To address this, Gulrajani et al. (2017) introduce a gradient penalty that improves stability by penalizing large-norm gradients, at the cost of longer computation; this variant is known as WGAN-GP.

- **Conditional Tabular GAN (CTGAN)** adds two tabular-specific components on top of a WGAN-GP objective: mode-specific normalization, which fits a Gaussian mixture to each continuous column to represent multimodal marginals accurately, and a conditional generation vector, which counteracts severe under-representation of minority categories during training (Xu et al., 2019).
- **Wasserstein GAN with Gradient Penalty (WGAN-GP)** is included as the general-purpose adversarial baseline without tabular-specific adaptation, which isolates how much of CTGAN’s performance is attributable to its tabular-specific components versus the shared adversarial training mechanism.

### 3.3. Evaluation Framework

We evaluate synthetic data across three primary dimensions: resemblance, utility, and privacy, following Hernadez et al. (2023). We additionally characterize the synthesis instability via variance decomposition from the two-stage imputation-synthesis design.

**Resemblance** is evaluated univariately and multivariately.

*Univariate* resemblance analysis (URA) compares variable-by-variable marginal distributions using three Goodness-of-fit tests (Student’s *t*-test, Mann-Whitney *U*-test, and Kolmogorov-Smirnov test) to assess statistical distinguishability, and marginal Wasserstein distance to quantify the magnitude of discrepancy.

*Multivariate* resemblance analysis (MRA) compares the Spearman rank correlation matrix of real and synthetic data, summarized by two metrics. The first *MRA score* is the proportion of upper-triangular pairwise correlations whose absolute difference from the real data falls below 0.1 (Hernadez et al., 2023), ranging from 0 to 1, with higher values indicating better preservation of the pairwise dependence structure. Secondly, the *Frobenius norm*of the upper-triangular absolute correlation matrix **C** = (*c*_*jj*′_) ∈ ℝ^*p×p*^,

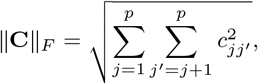

measures the overall magnitude of pairwise correlations. A smaller discrepancy between the synthetic and real Frobenius norm indicates better-preserved aggregate dependence strength.

In addition, *Data Labeling Analysis* (DLA) is used to assess semantic indistinguishability by training machine learning classifiers to discriminate real from synthetic data. We considered five classifiers:

1. Random Forest (RF), the ensemble of decision trees that aggregates predictions via majority vote to reduce variance;
2. *K*-Nearest Neighbors classifier (KNN), where class labels are assigned based on the majority class among the *k* closest training observations;
3. Decision Tree (DT), the recursive partitioning classifier that produces interpretable axis-aligned decision rules;
4. Support Vector Machine (SVM) with a linear kernel, which seeks the maximum-margin separating hyperplane in the feature space;
5. Multilayer Perceptron (MLP), the feedforward neural network with fully connected hidden layers trained by backpropagation.

Accuracy, precision, recall, and F1-score near chance level (0.5) indicate high fidelity, since no reliable distinction can be drawn between real and synthetic data.

**Utility** is assessed via the Train-on-*Real*-Test-on-Real (TRTR) and Train-on-*Synthetic*-Test-on-Real (TSTR) paradigm with the prespecified classifiers above (Hernadez et al., 2023). Models trained on real and synthetic data are each evaluated on the same held-out real test set.

For *supervised* learning, smaller absolute differences between TRTR and TSTR values of accuracy, precision, recall, and F1-score indicate that synthetic data supports supervised learning comparably well to real data, i.e., with high supervised learning utility.

For *unsupervised* learning, we evaluated clustering performance using group-based multi-trajectory modeling (GBMT), specialized for longitudinal data (Nagin et al., 2018). GBMT was fit separately on the real and synthetic training sets, and the resulting models were used to assign trajectory groups to the held-out test set. These assignments were compared against the true group labels using the Adjusted Rand Index (ARI), yielding ARI_TRTR_ and ARI_TSTR_, whose closer agreement indicates better preservation of the dataset’s unsupervised structure by the synthetic data (Rand, 1971; Hubert and Arabie, 1985).

**Privacy** is evaluated via a simulated *Membership Inference Attack* (MIA), in which a hypothetical attacker attempts to determine whether a real record was used to train the generator. Before the attack, all continuous variables were discretized into five quantile-based bins and encoded as integer categories. Record dissimilarity was then measured using Hamming distance, defined as the proportion of attributes with different category values. For example, bin 3 versus bin 3 counts as a match, whereas bin 3 versus bin 5 counts as a mismatch. At each of four decision thresholds (0.1, 0.2, 0.3, and 0.4), a real record was classified as a training member if at least one synthetic record fell within the specified Hamming distance threshold. Smaller thresholds therefore impose stricter matching criteria.

To simulate varying levels of attacker knowledge, we constructed an *evaluation pool* containing equal numbers of records from the training and held-out test sets. For each attacker’s knowledge level (shown as the x-axis of **Figure 6**), the attacker was assumed to know the corresponding proportion of records randomly sampled from this pool. Each sampled record’s true membership label was then compared with the label predicted by the Hamming-distance rule, yielding a standard binary classification problem.

We summarized attack performance using accuracy and precision. Because the evaluation pool was balanced, accuracy near 50% indicates chance-level membership classification and therefore stronger privacy preservation, whereas accuracy substantially above or below 50% indicates greater information leakage. Precision provided a complementary measure, quantifying the proportion of records flagged as members that were truly drawn from the training set. Elevated precision was interpreted as evidence of greater re-identification risk: higher precision indicates that records flagged as members are more often true training records, whereas low precision suggests weaker membership identification but should be interpreted in light of the number of positive predictions (Hernadez et al., 2023).

**Variability** was assessed using the descriptive variance decomposition described in Section 2.4. For each method, we computed the grand mean Wasserstein distance 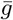, the bias-corrected between-imputation variance 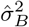, and the within-imputation synthesis variance 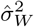 across the *M* ×*L* = 50 dataset ensemble (*M* = 5 imputations and *L* = 10 syntheses per imputation).

We summarize the evaluation framework, including the methods assessed under each evaluation criterion, in **Table 1**.

**Table 1.**
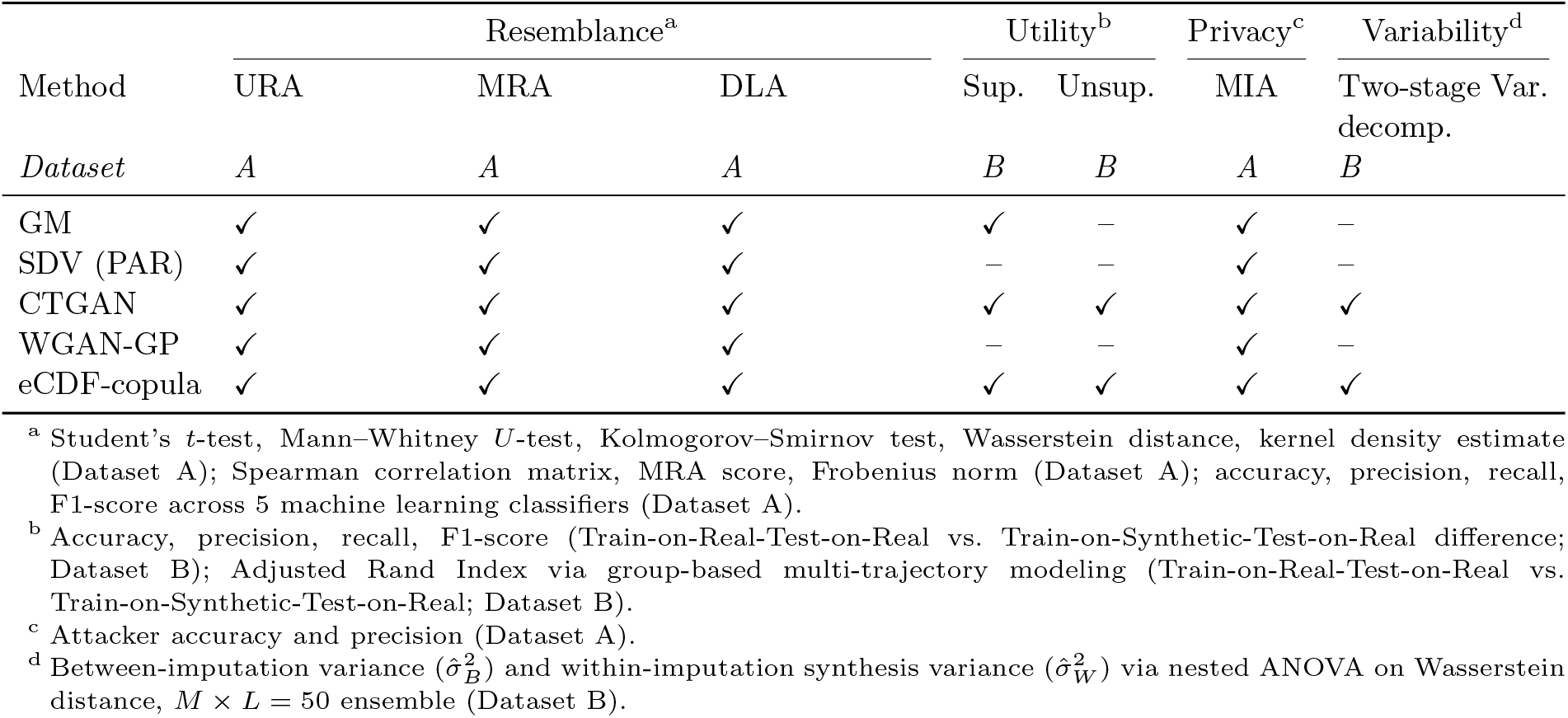
Overview of the evaluation framework for five synthetic data generation methods. ✓ indicates the method was included in the corresponding evaluation; – indicates it was not. Dataset A: REMBRANDT (*n* = 120); Dataset B: CHAP (*n* = 3,612). Abbreviations: GM, Gaussian multivariate; SDV, synthetic data vault; PAR, probabilistic autoregressive model; CTGAN, conditional tabular generative adversarial network; WGAN-GP, Wasserstein generative adversarial network with gradient penalty; URA, Univariate Resemblance Analysis; MRA, Multivariate Resemblance Analysis; DLA, Data Labeling Analysis; MIA, Membership Inference Attack; Sup., supervised; Unsup., unsupervised; Var. decomp., variance decomposition

| Method | Resemblance<sup>a</sup> |  |  | Utility<sup>b</sup> |  | Privacy<sup>c</sup> | Variability<sup>d</sup> |
| --- | --- | --- | --- | --- | --- | --- | --- |
|  | URA | MRA | DLA | Sup. | Unsup. | MIA | Two-stage Var. decomp. |
| *Dataset* | *A* | *A* | *A* | *B* | *B* | *A* | *B* |
| GM | ✓ | ✓ | ✓ | ✓ | – | ✓ | – |
| SDV (PAR) | ✓ | ✓ | ✓ | – | – | ✓ | – |
| CTGAN | ✓ | ✓ | ✓ | ✓ | ✓ | ✓ | ✓ |
| WGAN-GP | ✓ | ✓ | ✓ | – | – | ✓ | – |
| eCDF-copula | ✓ | ✓ | ✓ | ✓ | ✓ | ✓ | ✓ |

<sup>a</sup> Student’s $t$ -test, Mann–Whitney $U$ -test, Kolmogorov–Smirnov test, Wasserstein distance, kernel density estimate (Dataset A); Spearman correlation matrix, MRA score, Frobenius norm (Dataset A); accuracy, precision, recall, F1-score across 5 machine learning classifiers (Dataset A).
<sup>b</sup> Accuracy, precision, recall, F1-score (Train-on-Real-Test-on-Real vs. Train-on-Synthetic-Test-on-Real difference; Dataset B); Adjusted Rand Index via group-based multi-trajectory modeling (Train-on-Real-Test-on-Real vs. Train-on-Synthetic-Test-on-Real; Dataset B).
<sup>c</sup> Attacker accuracy and precision (Dataset A).
<sup>d</sup> Between-imputation variance ( $\hat{\sigma}_B^2$ ) and within-imputation synthesis variance ( $\hat{\sigma}_W^2$ ) via nested ANOVA on Wasserstein distance, $M \times L = 50$ ensemble (Dataset B).

## 4. Results

We present results in the four dimensions. For resemblance and privacy evaluations (Sections 4.1 and 4.3), we focused on Dataset A (REMBRANDT, *n* = 120), whose compact size and repeated-measures structure pose a challenging scenario for assessing marginal and multivariate fidelity, and a stringent setting for privacy, where smaller *n* affords less inherent anonymity. Utility and variability evaluations (Sections 4.2 and 4.4) are reported based on Dataset B (CHAP, *n* = 3, 612), as its larger sample size and binary group indicator provide the stability required for TRTR versus TSTR comparisons across supervised and unsupervised learning tasks.

### 4.1. Resemblance

As introduced in Section 3.3, resemblance was assessed through univariate analysis, multivariate analysis, and data labeling analysis.

#### 4.1.1. Univariate Resemblance Analysis

We evaluated univariate resemblance of the five synthetic approaches across 28 longitudinal variables (MADRS and HARS composite scores at 14 visits) from the Dataset A.

**Goodness-of-fit tests**. The null hypothesis is marginal distributional equivalence between real and synthetic data. Across all three tests (Student’s *t*-test, Mann-Whitney *U*-test, and Kolmogorov-Smirnov test), the proposed eCDF-copula consistently failed to reject the null, supporting its distributional fidelity to the real data (predominantly green in **Figure 1**). In contrast, near-zero *p*-values across almost all variables in WGAN-GP and CTGAN indicate substantial deviation from the real marginals, consistent with their lack of explicit mechanisms for preserving temporal dependence in longitudinal data and the data-hungry nature of adversarial training (Miletic and Sariyar, 2025; Xu et al., 2019). GM and SDV showed intermediate performance, exceeding the 0.05 threshold for some but not all variables.

**Figure 1.**
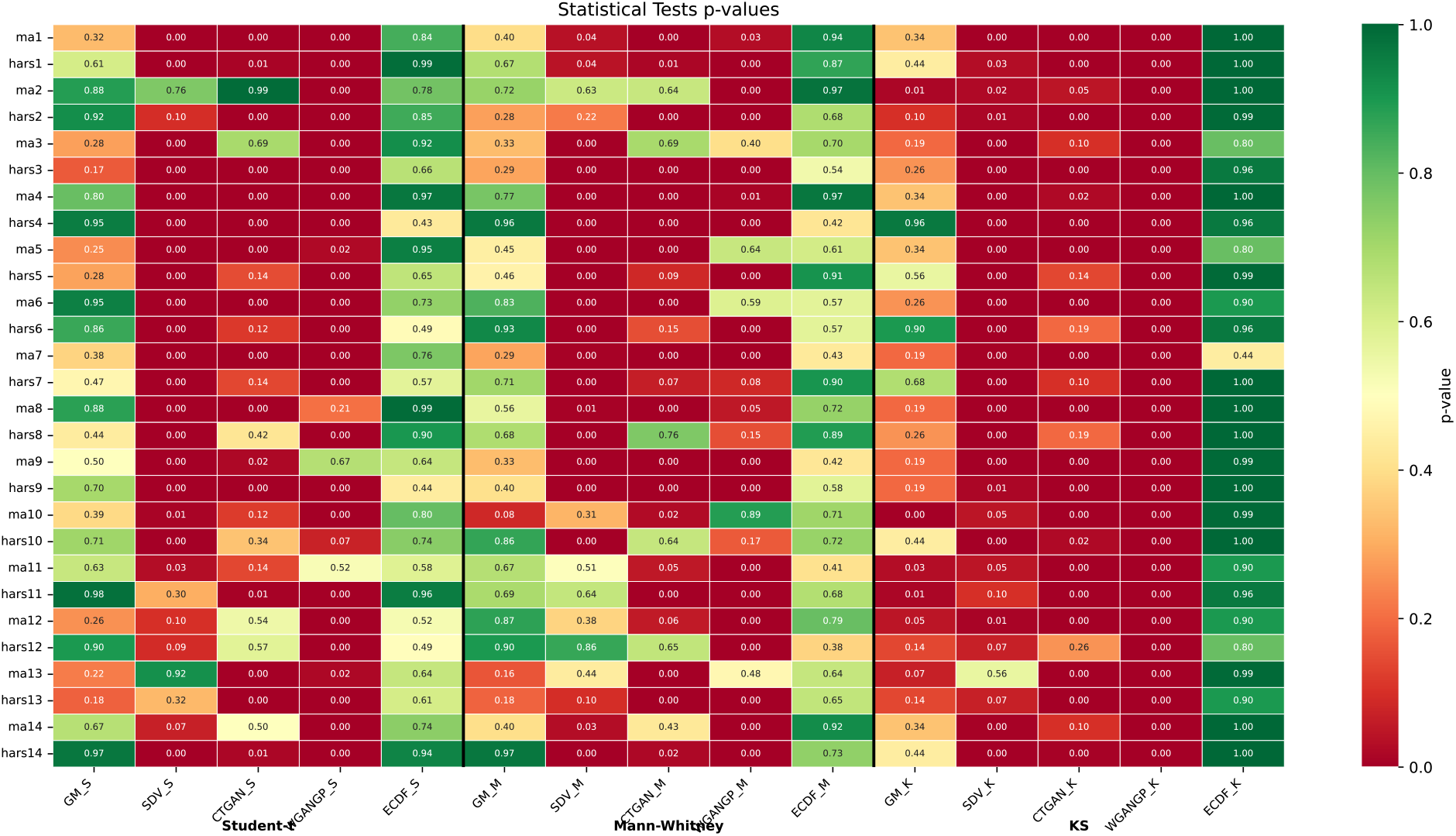
Heatmap of *p*-values from three univariate statistical tests (Student’s *t*-test, Mann–Whitney *U*-test, and Kolmogorov–Smirnov test) comparing real and synthetic data distributions across 28 longitudinal variables (MADRS and HARS scores at 14 visits). Higher *p*-values (green) indicate failure to reject the null hypothesis of distributional equivalence. Columns correspond to five synthetic data generation methods: GM, SDV, CTGAN, WGAN-GP, and the proposed eCDF-copula approach.

**Marginal Wasserstein distance and density estimates**. In **Table 2**, we report the mean (standard deviation) Wasserstein distance between real and synthetic marginal distributions. eCDF-copula achieved the lowest mean distance at 0.034, followed by SDV (0.066), GM (0.098), CTGAN (0.103), and WGAN-GP (0.172). Further, visual inspection using kernel density estimates for each variable (see **Supplementary Figure S1**) confirmed this resemblance. Interestingly, WGAN-GP generated markedly over-concentrated distributions, with near-degenerate spikes around certain values. This mode collapse behavior has been previously observed in GANs (Alqahtani et al., 2021), where the generator converges to a narrow region when adversarial training is destabilized by insufficient sample size (Gulrajani et al., 2017). Additionally, the miscalibrated distributions from CTGAN reflect a related failure mode driven by the same moderate-sample-size constraint common in clinical studies, distinct from the large-scale data regimes for which these models were originally developed.

**Table 2.** Univariate and multivariate resemblance metrics for five synthetic data generation methods, computed across 28 longitudinal variables (MADRS and HARS scores at 14 visits). Arrows indicate the direction of improvement (↑ higher is better; ↓ lower is better). Wasserstein: mean (SD) Wasserstein distance between real and synthetic marginal distributions (↓). Multivariate Resemblance Analysis (MRA) Score: proportion of pairwise correlations within 0.1 of the real data (↑; range 0–1). |Δ_*F*_ |: absolute difference between the synthetic and real data Frobenius norms (∥**C**_real_∥_*F*_ = 14.45; ↓). Bold indicates the best synthetic method in each column.

| Method | Wasserstein (↓) | MRA Score (↑) | $\vert \Delta_F\vert $ (↓) |
| --- | --- | --- | --- |
| GM | 0.098 (0.038) | 0.59 | 0.72 |
| SDV | 0.066 (0.028) | 0.07 | 6.65 |
| CTGAN | 0.103 (0.052) | 0.03 | 8.31 |
| WGAN-GP | 0.172 (0.084) | 0.35 | 1.24 |
| eCDF-copula (proposed) | **0.034 (0.024)** | **0.61** | **0.62** |

#### 4.1.2. Multivariate Resemblance Analysis

To evaluate the multivariate dependence structure, we compared Spearman rank correlation matrices between real and synthetic data, the MRA scores, and the discrepancy between synthetic and real Frobenius norms (∥**C**_real_∥_*F*_ = 14.45).

The real data showed a block correlation pattern reflecting correlation among repeated measures over time. eCDF-copula not only reproduced this structure visually (**Figure 2**), but also attained the highest MRA (Multivariate Resemblance Analysis) score (0.61) and yielded the Frobenius norm closest to real (∥**C**_syn_∥_*F*_ = 13.83, |Δ_*F*_ | = 0.62, **Table 2**). GM performed comparably (MRA = 0.59, |Δ_*F*_ | = 0.72). SDV and CTGAN, however, substantially underestimated dependence, with near-zero MRA scores (0.07, 0.03) and Frobenius norms far below the real value (|Δ_*F*_ |: 6.65, 8.31). WGAN-GP inflated overall correlation magnitude (|Δ_*F*_ | = 1.24) while achieving only a moderate MRA score (0.35), indicating it overstated dependence strength without faithfully reproducing the pairwise pattern.

**Figure 2.**
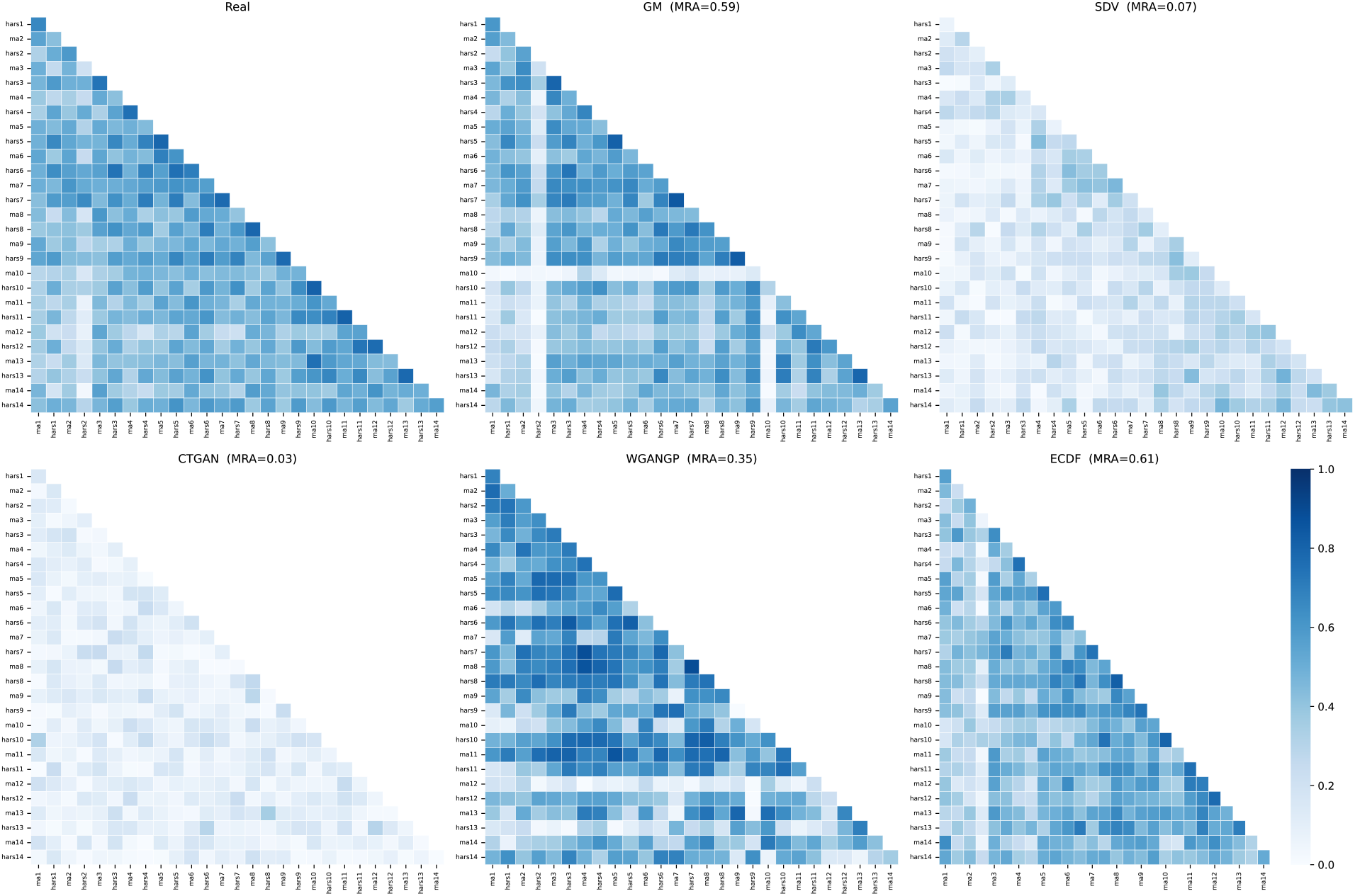
Spearman rank correlation matrices for the real data and five synthetic data generation methods, computed across 28 longitudinal variables (MADRS and HARS scores at 14 visits). The MRA score (the proportion of pairwise correlations within 0.1 of the real data) and the Frobenius norm are reported for each method. The real data Frobenius norm is 14.45.

#### 4.1.3 Data Labeling Analysis

For each method, we trained the five classifiers to distinguish real from synthetic data. **Figure 3** summarizes the resulting accuracy, precision, recall, and F1-score, with each box reflecting the spread of a metric across the five classifiers. Performance near chance level (0.5) indicates high fidelity, suggesting that the classifiers cannot reliably tell synthetic data from real.

**Figure 3.**
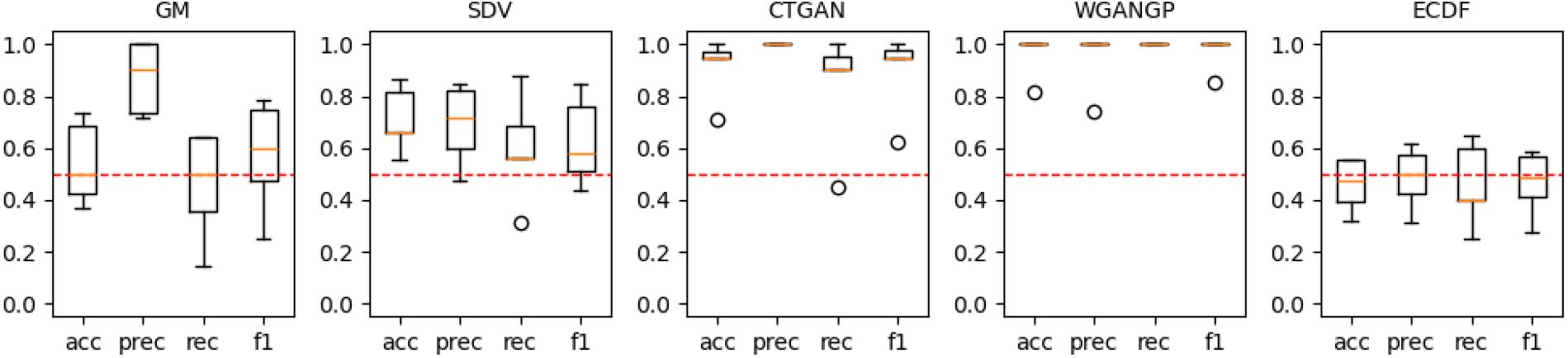
Box plots of classification performance metrics (accuracy, precision, recall, and F1-score) from the Data Labeling Analysis (DLA). A panel of five machine learning classifiers (Random Forest, K-Nearest Neighbors, Decision Tree, SVM, and MLP) was trained to distinguish real from synthetic records; each box summarizes the spread of the metric across these five classifiers. Values near 0.5 indicate chance-level discrimination, reflecting high semantic indistinguishability of the synthetic data. Results are shown for GM, SDV, CTGAN, WGAN-GP, and the proposed eCDF-copula method. The dashed red line marks the 0.5 reference.

The proposed eCDF-copula method achieved the strongest semantic indistinguishability: it was the only method whose interquartile range consistently covered 0.5 across all four metrics, while also exhibiting relatively low variability across classifiers. In contrast, CTGAN and WGAN-GP produced near-ceiling metrics (median approaching 1.0), indicating that they are trivially distinguishable from real tabular data. GM and SDV showed intermediate performance but with considerable variability across classifiers.

### 4.2. Utility

Utility was evaluated across supervised classification and unsupervised clustering under Train-on-Real-Test-on-Real (TRTR) and Train-on-Synthetic-Test-on-Real (TSTR). We focus this section on Dataset B (*n* = 3,612), with a larger sample size to stabilize estimation. Parallel results for Dataset A (*n* = 120), a more challenging small-sample regime, are reported in the **Supplementary Material**.

#### 4.2.1. Supervised Learning

We trained the five classifiers (RF, KNN, DT, SVM, and MLP) using real and synthetic data, respectively. **Figure 4** shows the absolute differences in four classification metrics (accuracy, precision, recall, and F1-score) between TRTR and TSTR across these classifiers; smaller differences indicate that the synthetic data more faithfully substitutes for real data in these learning tasks.

**Figure 4.**
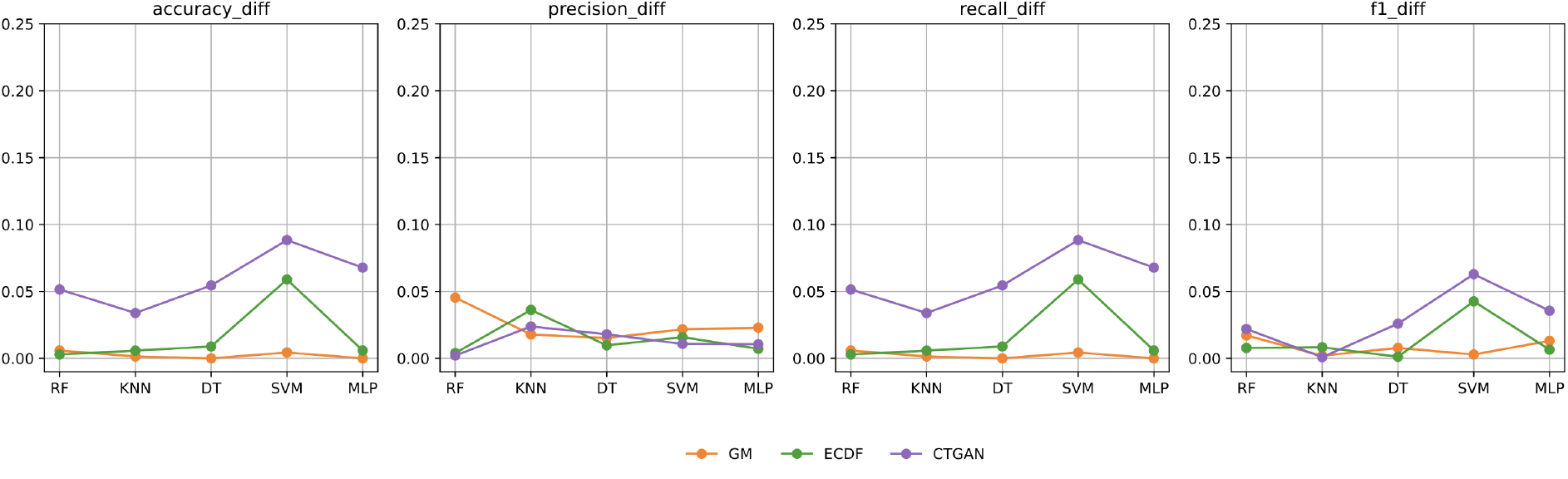
Absolute differences in classification performance metrics (accuracy, precision, recall, F1-score) between Train-on-Real-Test-on-Real (TRTR) and Train-on-Synthetic-Test-on-Real (TSTR) across five classifiers (RF, KNN, DT, SVM, MLP) for the dataset B. Smaller values indicate higher utility of the synthetic data for supervised learning.

Among the synthetic data methods, GM showed the lowest TRTR-TSTR differences (all metrics below 0.05), indicating that classifiers trained on GM-synthesized data achieved comparable utility to those trained on real data. The proposed eCDF-copula method maintained low differences for RF, KNN, DT, and MLP, but exhibited a modest spike at SVM. CTGAN showed the largest discrepancies among all synthetic methods, peaking at SVM across multiple metrics.

Across classifiers, the support vector machine (SVM) consistently showed the largest discrepancies regardless of the synthesis method. As noted in prior studies (Patki et al., 2016; Zhang et al., 2022; Hernadez et al., 2023), margin-based classifiers tend to be less robust to distributional discrepancies between training and test sets than other methods, where small shifts in the marginal distributions or covariance structure of synthetic data can alter the position of the separating hyperplane and thereby inflate TSTR metric differences relative to TRTR.

#### 4.2.2. Unsupervised Learning

We next evaluated unsupervised utility for clustering multivariate longitudinal trajectories using group-based multi-trajectory modeling (GBMT), comparing clustering performance via the Adjusted Rand Index (ARI). Under TRTR, the GBMT was fitted on real data and applied to held-out real observations; under TSTR, the GBMT was fitted on synthetic data and applied to the same held-out observations. Closer agreement between ARI_TRTR_ and ARI_TSTR_ indicates that synthetic data better preserves the unsupervised utility of the original dataset.

Here, we focus on the comparison between eCDF-copula and CTGAN, since SDV and WGAN-GP generated severely imbalanced synthetic group labels, synthesizing predominantly one group, and are unsuitable for clustering evaluation. Results (see **Figure 5**) are reported across 50 replication datasets generated following the procedure described in Section 4.4.

**Figure 5.**
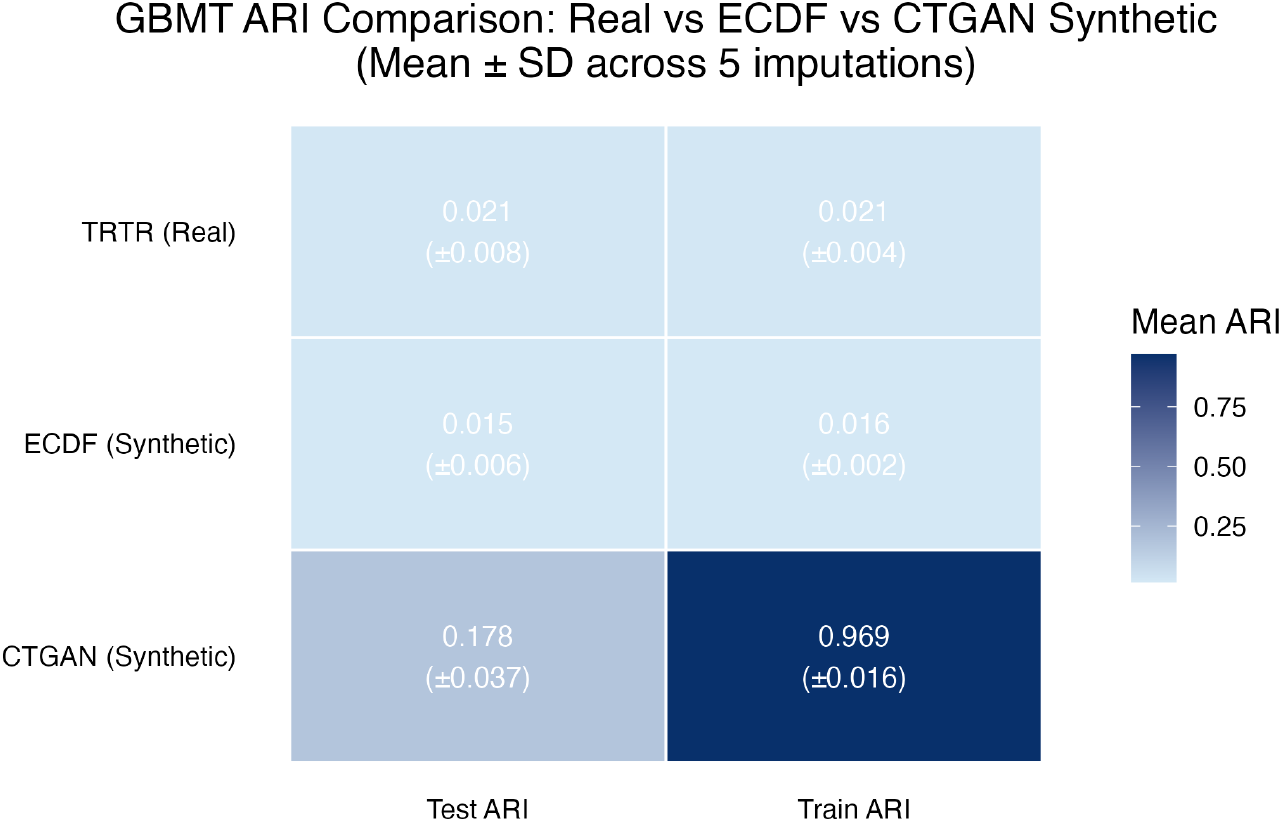
GBMT clustering utility comparison on CHAP dataset. Mean Adjusted Rand Index (ARI; s.d. in parentheses) across 5 multiple imputations under TRTR (real data) and TSTR (eCDF-copula and CTGAN synthetic data), evaluated on both training and test partitions. Values closer to the TRTR benchmark indicate higher clustering utility.

Under TRTR, the real data yielded a mean ARI_TRTR_ of 0.021 (s.d. 0.008) on the training set and 0.021 (s.d. 0.004) on the test set, reflecting the modest but consistent cluster separability present in the original trajectories. eCDF-copula closely replicated this pattern, with ARI_TSTR_ values of 0.015 (s.d. 0.006) on training and 0.016 (s.d. 0.002) on test, with differences from the real data remaining within the range of imputation-induced variability. CTGAN, however, exhibited severe overfitting: its ARI_TRTR_ on the training set was 0.969 (s.d. 0.016), indicating near-perfect separation of cluster labels in the synthetic training data, while its ARI_TSTR_ dropped to 0.178 (s.d. 0.037), still far exceeding the real-data benchmark and demonstrating that CTGAN-generated data does not generalize to held-out real observations.

Dataset A revealed a similar pattern: eCDF-copula showed moderate deviation between ARI_TRTR_ and ARI_TSTR_, while SDV exhibited the same overfitting behavior observed for CTGAN on Dataset B, with a high ARI_TRTR_ but low ARI_TSTR_ (see **Supplementary Figures S2 and S3**).

### 4.3. Privacy

Privacy was assessed using the simulated Membership Inference Attack described in Section 3.3, with attacker *accuracy* and *precision* computed across four Hamming-distance thresholds (0.1–0.4). Throughout, accuracy near 50% indicates chance-level membership classification and stronger privacy preservation, and higher precision indicates greater re-identification risk.

In terms of *accuracy*, almost all methods maintained chance-level attack performance across thresholds and attacker knowledge levels (**Figure 6**). As the Hamming-distance threshold was relaxed, however, *precision* differed more clearly across methods, though estimates were unstable when the number of records flagged as members was very small. Consequently, the precision of GM and eCDF-copula at thresholds of 0.1 and 0.2 jumped from 0 to 1 and is therefore not practically meaningful. CTGAN and SDV showed near-zero precision even at looser thresholds of 0.2 and 0.3, suggesting that records flagged as members were rarely true training records. Focusing instead on the more informative range, WGAN-GP at a threshold of 0.3 showed elevated and increasing precision relative to the other methods, indicating the highest privacy risk.

### 4.4. Variability

Variability was assessed via the decomposition described in Section 3.3, with between-imputation variance 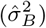 and within-imputation synthesis variance 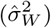 computed for eCDF-copula and CTGAN across the *M* × *L* = 50 dataset ensemble (*M* = 5 imputations × *L* = 10 syntheses).

Overall, eCDF-copula achieved a substantially lower grand mean Wasserstein distance than CTGAN (0.023 vs. 0.073), indicating higher marginal fidelity. The variance decomposition revealed distinct uncertainty profiles (**Figure 7**; all variance values ×10^*−*6^). eCDF-copula showed lower between-imputation variance than CTGAN (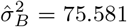vs. 173.488), suggesting that its marginal fidelity was less sensitive to which imputed dataset was used for training. However, eCDF-copula showed greater within-imputation synthesis variance (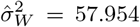 vs. 9.074), reflecting additional stochasticity from the copula sampling step. In contrast, CTGAN showed lower conditional synthesis variability but greater sensitivity to the imputed training dataset. Taken together, eCDF-copula had lower total variability, 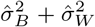 than CTGAN, indicating that its improved marginal fidelity was not achieved at the cost of greater overall instability.

**Figure 6.**
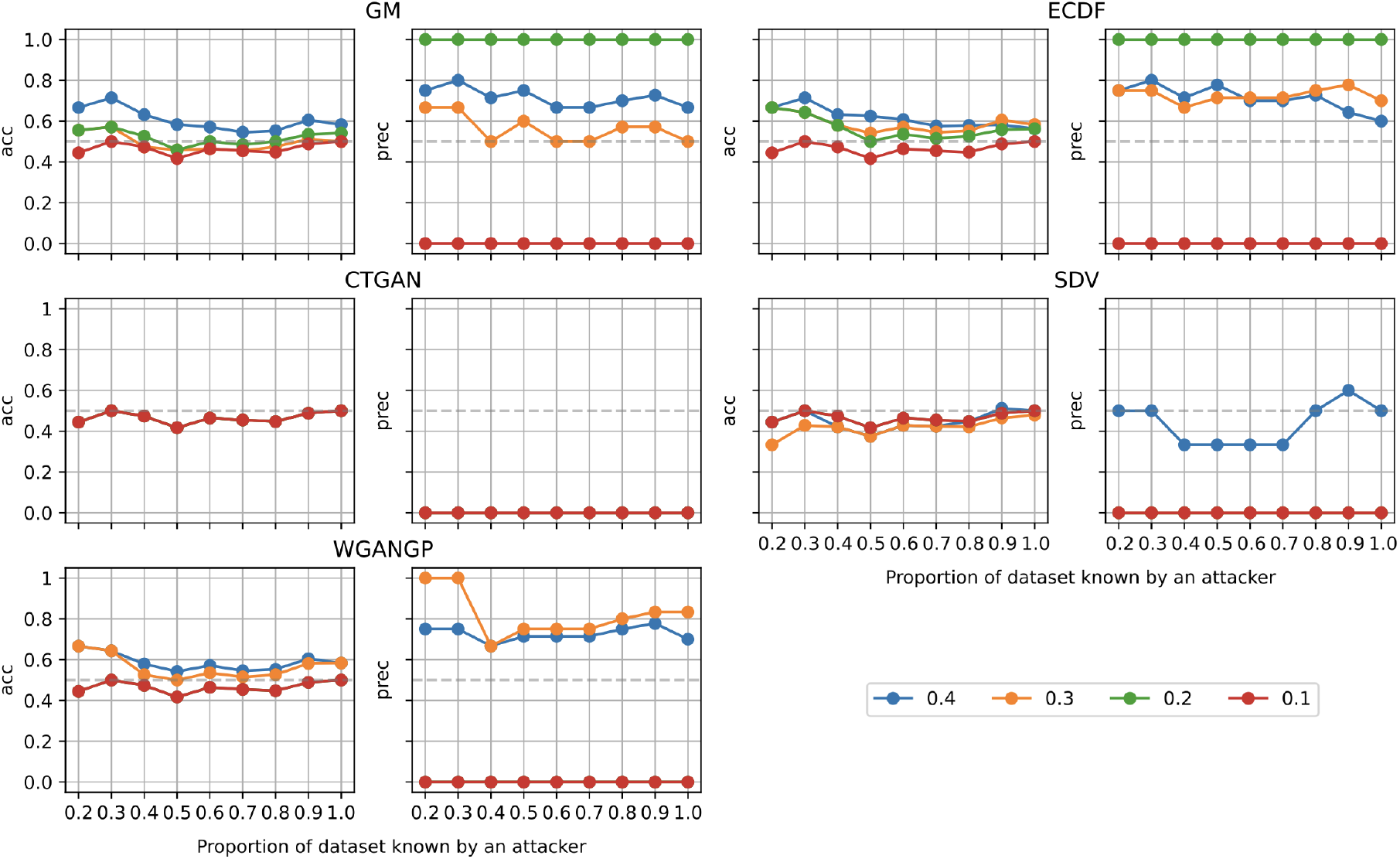
Simulated Membership Inference Attack (MIA) results for five synthetic data generation methods on the REMBRANDT dataset A. Each panel shows attacker accuracy (acc, left) and precision (prec, right) as a function of the proportion of the real dataset known to the attacker (x-axis). Line colors indicate Hamming distance thresholds (0.1, 0.2, 0.3, 0.4); a smaller threshold requires a closer match between synthetic and real records for the attacker to claim identification. The dashed grey line marks the chance level (50%). Accuracy near 50% indicates chance-level membership classification and stronger privacy preservation, and higher precision indicates greater re-identification risk.

**Figure 7.**
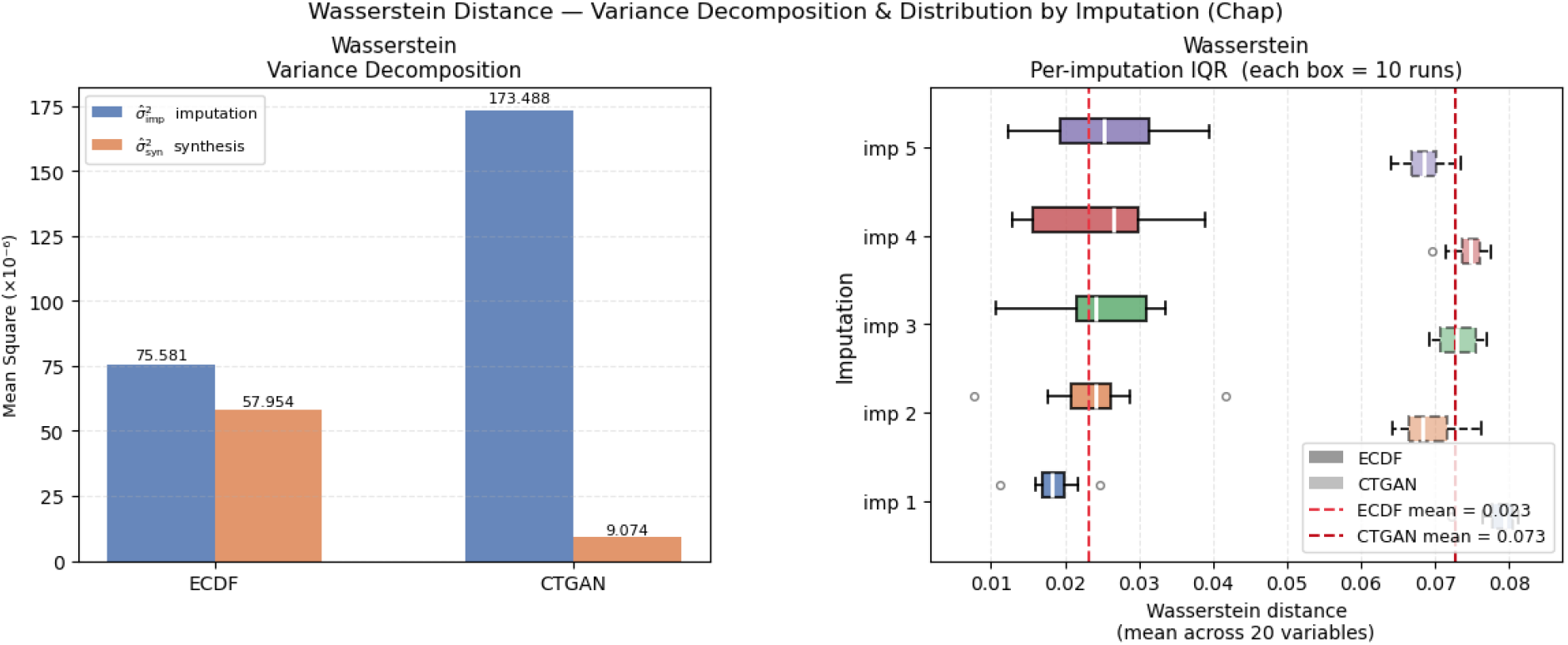
Variance decomposition and per-imputation distribution of mean Wasserstein distance across 20 variables for eCDF-copula and CTGAN, based on the *M* × *L* = 50 ensemble (5 imputations × 10 syntheses per method) on Dataset B. *Left:* Between-imputation variance (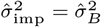, blue) and pooled within-imputation synthesis variance 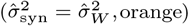 estimated via nested ANOVA mean squares (×10^*−*6^). *Right:* Per-imputation interquartile ranges of mean Wasserstein distance; each box summarizes *L* = 10 synthesis replicates within a single imputation replicate (imp 1–5). Red dashed vertical lines indicate the per-method grand mean Wasserstein distance (eCDF-copula: 0.023; CTGAN: 0.073).

## 5. Discussion

We proposed a statistically grounded eCDF-copula method for synthetic data generation, extending the cross-sectional construction to longitudinal settings. We evaluated the proposed method against four established approaches, Gaussian multivariate (GM), probabilistic autoregressive via Synthetic Data Vault (SDV), and two generative adversarial network (GAN)-based approaches, CTGAN and WGAN-GP, on two longitudinal datasets spanning small and large sample sizes (*n* = 120 and *n* = 3,612).

Across the primary resemblance, utility, and privacy dimensions, eCDF-copula performed consistently well relative to the competing methods. For *resemblance*, it achieved the lowest univariate Wasserstein distance, preserved multivariate dependence most effectively, and generated synthetic data that was hardest to distinguish from real data under data labeling analysis. For *utility*, synthetic data from eCDF-copula yielded nearly identical downstream results to those trained on real data for both supervised and unsupervised learning tasks. Moreover, unlike CTGAN, whose synthetic data overfit to the training set, eCDF-copula reproduced the real clustering structure more faithfully. Finally, for *privacy*, membership inference attacks revealed threshold-dependent privacy risk: CTGAN, SDV, and eCDF-copula maintained reasonable attacker performance, while WGAN-GP showed elevated re-identification risk at looser matching criteria.

Further, *variability* across repeated syntheses offers additional insight. eCDF-copula showed lower between-imputation variance than CTGAN but higher within-imputation synthesis variance, reflecting the added stochasticity of copula sampling. Even so, its total variability was lower than CTGAN’s, so its higher fidelity did not come at the cost of greater overall instability. Whether this trade-off generalizes to other datasets and synthesis settings is a direction for future work.

Beyond this trade-off, eCDF-copula’s overall superior performance can be explained by limitations of each competing approach. Both CTGAN and WGAN-GP are designed for cross-sectional tabular data (Xu et al., 2019; Alqahtani et al., 2021) and, when applied to longitudinal data, treat the stacked visit-variable matrix as an ordinary flat table, thereby discarding the temporal structure that contributed to CTGAN’s near-zero MRA score (0.03) and WGAN-GP’s degraded multivariate fidelity (MRA = 0.35). More importantly, adversarial training requires sufficiently diverse real-data gradients to guide the generator. With only limited subjects in small-to-moderate datasets, the discriminator rapidly memorizes the training data, leading to near-degenerate mode collapse in WGAN-GP and spurious multimodality in CTGAN (Gulrajani et al., 2017; Heine et al., 2023). eCDF-copula’s advantage over GM, by contrast, stems mainly from its treatment of marginal distributions: GM imposes parametric families on each variable (Hernadez et al., 2023), introducing systematic bias under the skewness and bounded support characteristic of clinical measures, whereas eCDF-copula’s nonparametric marginals preserve observed distributional features without parametric assumptions (Restrepo et al., 2023; Ben Hassine and Mili, 2025). In contrast, the poor multivariate resemblance of SDV (MRA = 0.07) is due to its autoregressive architecture which captures within-subject temporal evolution but not simultaneous cross-variable correlations at each visit (Miletic and Sariyar, 2025).

Despite these strengths, our proposed method has some limitations. First, eCDF-copula’s higher fidelity came with lower ensemble stability across replicates than CTGAN’s within-imputation behavior, though its total variability remained lower overall. Second, our implementation performs synthesis separately within each treatment group, which improves fidelity to group-specific structure but assumes group labels are known and available, a condition typically met by analysts with legitimate access to clinical data but not guaranteed in all deployment settings. Third, privacy was assessed using a simplified membership inference attack rather than a formal or worst-case privacy guarantee, and near-chance attacker performance under this specific attack should not be interpreted as evidence of resistance to other, potentially stronger, adversarial strategies. Future work includes developing variance-reduction strategies within the copula synthesis framework, such as importance sampling or control variates, to reduce replicate variability without sacrificing fidelity, as well as evaluating privacy under a broader range of attack models to more rigorously characterize re-identification risk.

In conclusion, eCDF-copula provides a practical solution for generating longitudinal clinical data, with strong fidelity and downstream utility across diverse evaluation metrics. It facilitates secure data sharing in biomedical research and paves the way for broader use of nonparametric copula-based synthesis in sensitive clinical studies.

## Acknowledgements

The authors gratefully acknowledge the investigators of the REMBRANDT and CHAP studies for providing access to the datasets used in this research.

## Data availability

The REMBRANDT and CHAP datasets analyzed in this study were obtained from the studies described in Taylor et al. (2024) and Tita et al. (2022), respectively, and are not publicly available due to privacy restrictions. Access to the data may be requested from the corresponding authors of the original studies. The code implementing the ECDF-copula method, along with scripts to reproduce the evaluation results and analyses presented in this paper, is available at https://github.com/hanchangcai2002/ecdf-copula. The repository is currently private and will be made public upon acceptance of this article.

