## Supplementary Figures Sx for "Synthetic Longitudinal Tabular Data Generation via Copula"

#### Supplementary Materials

Hanchang Cai,[1] Wenshan Yu,[2] Ruijin Lu,[3] Ishanu Chattopadhyay,[4] Xinlian Zhang[1] and Jinyuan Liu[5,*]

[1]Division of Biostatistics and Bioinformatics, Herbert Wertheim School of Public Health and Human Longevity Science, University of California San Diego, 9500 Gilman Drive, La Jolla, CA 92093, United States
[2]Department of Population Health Sciences, School of Public Health, Georgia State University, 140 Decatur Street, Atlanta, GA 30303, United States
[3]Institute for Informatics, Data Science & Biostatistics, Washington University in St. Louis School of Medicine, 660 S. Euclid Ave., St. Louis, MO 63110, United States
[4]Institute for Biomedical Informatics, Department of Internal Medicine, University of Kentucky College of Medicine, 800 Rose Street, Lexington, KY 40536,United States
[5]Departments of Biostatistics, Vanderbilt University Medical Center, 2525 West End Avenue, Suite 1100, Nashville, TN 37203, United States

#### Abstract

**Synthetic data generation is increasingly used to enable data sharing and secondary analysis while protecting participant privacy, particularly for longitudinal tabular health data, where repeated measures per subject create within-subject dependence that most synthetic data methods are not designed to preserve. Existing generative methods, particularly generative adversarial network (GAN)-based approaches, can model complex distributions, but their estimated dependence structures are often difficult to interpret and their performance may be unstable or prone to overfitting in modestly sized datasets. Here we show that eCDF-copula, a statistically rooted approach using the empirical cumulative distribution function (eCDF) and copula modeling, preserves within- and between-visit dependence structure. To handle pervasive missing data, we propose a two-stage strategy combining multiple imputation with copula-based synthesis, enabling a variance decomposition that quantifies replication variability across methods. We benchmarked the proposed approach against four established methods on two longitudinal clinical datasets spanning markedly different sample sizes ($n = 120$ vs. $n = 3,612$). eCDF-copula achieved resemblance and utility exceeding those of state-of-the-art synthetic data methods, while maintaining comparable privacy.**

**Keywords:** generative adversarial network (GAN); missing data; multiple imputation; synthesis variability; synthetic data evaluation; temporal correlation in longitudinal data

### Supplementary Figures

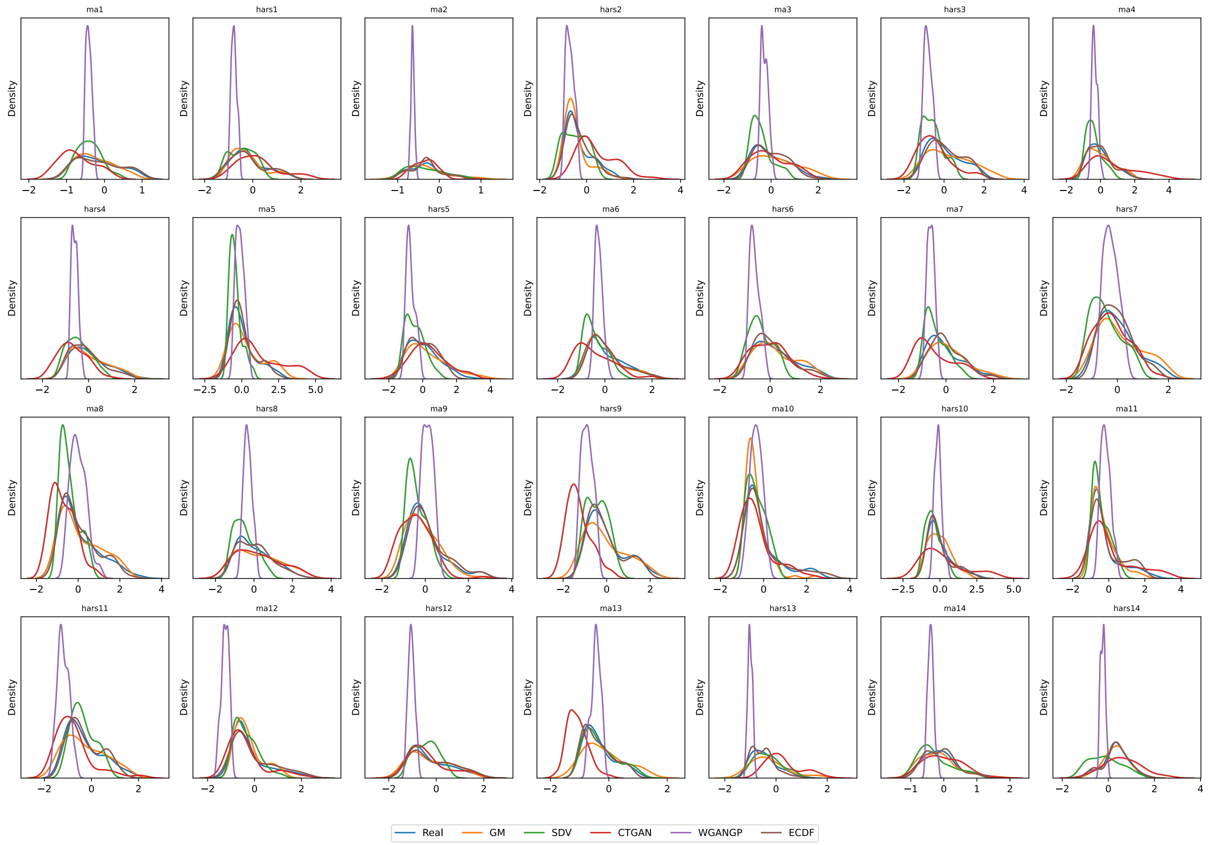


**Supplementary Figure S1.** Kernel density estimates of real and synthetic data distributions for each of the 28 longitudinal variables. Each panel displays the empirical density of the real data alongside those generated by GM, SDV, CTGAN, WGAN-GP, and the proposed ECDF-copula method.

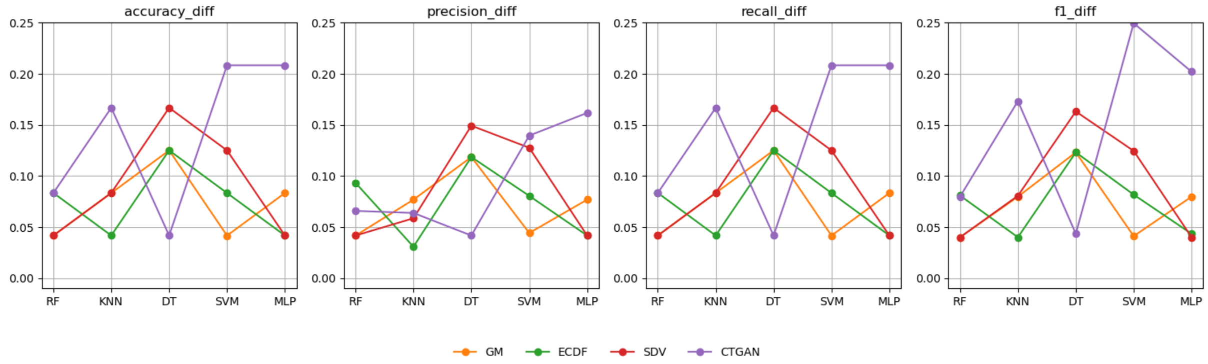


**Supplementary Figure S2.** Absolute differences in classification performance metrics (accuracy, precision, recall, and F1-score) between Train-on-Real-Test-on-Real (TRTR) and Train-on-Synthetic-Test-on-Real (TSTR) across five classifiers (RF, KNN, DT, SVM, and MLP) for the REMBRANDT dataset. Smaller values indicate higher utility of the synthetic data for supervised learning.

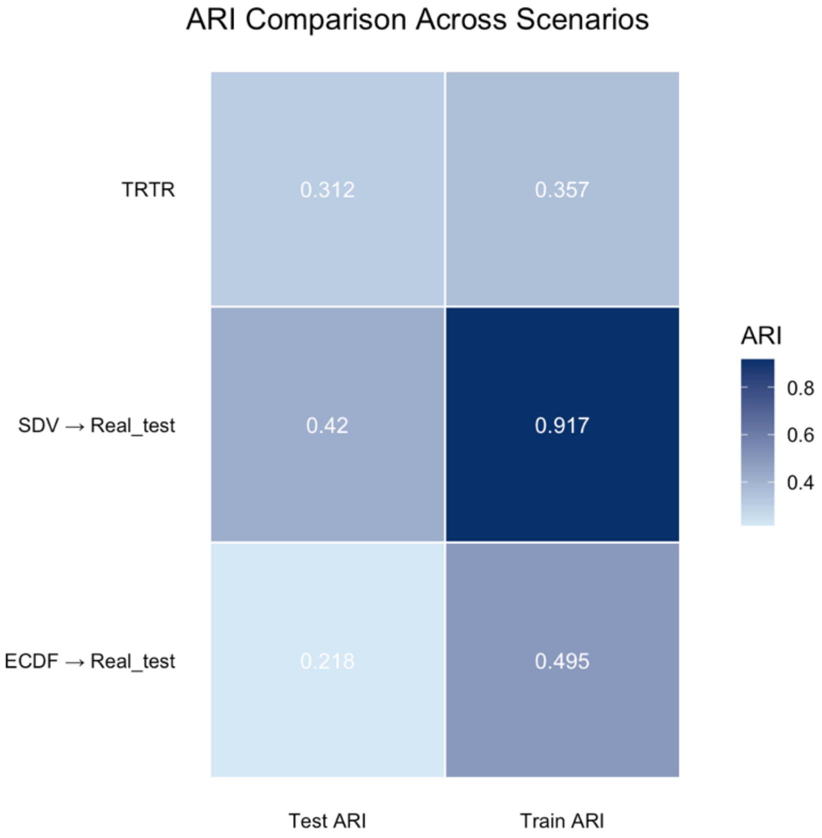


**Supplementary Figure S3.** GBMT clustering utility comparison on the REMBRANDT dataset. Adjusted Rand Index (ARI) under Train-on-Real-Test-on-Real (TRTR) and Train-on-Synthetic-Test-on-Real (TSTR; SDV and ECDF-copula synthetic data), evaluated on both training and test partitions. Values closer to the TRTR benchmark indicate higher clustering utility.
